# Mechanism of allosteric activation of human mRNA cap methyltransferase (RNMT) by RAM: Insights from accelerated molecular dynamics simulations

**DOI:** 10.1101/558502

**Authors:** Juan A. Bueren-Calabuig, Marcus Bage, Victoria H. Cowling, Andrei V. Pisliakov

## Abstract

The RNA guanine-7 methyltransferase (RNMT) in complex with RNMT-Activating Miniprotein (RAM) catalyses the formation of a N7-methylated guanosine cap structure on the 5’ end of nascent RNA polymerase II transcripts. The mRNA cap protects the transcript from exonucleases and recruits cap-binding complexes that mediate RNA processing, export and translation. By using microsecond standard and accelerated molecular dynamics simulations, we provide for the first time a detailed molecular mechanism of allosteric regulation of RNMT by RAM. We show that RAM selects the RNMT active site conformations that are optimal for binding of substrates (AdoMet and the cap), thus enhancing their affinity. Furthermore, our results strongly suggest the likely scenario in which the cap binding promotes the subsequent AdoMet binding, consistent with the previously suggested cooperative binding model. By employing the dynamic network and community analyses, we revealed the underlying long-range allosteric networks and paths that are crucial for allosteric regulation by RAM. Our findings complement and explain previous experimental data on RNMT activity. Moreover, this study provides the most complete description of the cap and AdoMet binding poses and interactions within the enzyme’s active site. This information is critical for the drug discovery efforts that consider RNMT as a promising anti-cancer target.

## INTRODUCTION

Enzymatic modification of the 5’-end of eukaryotic messenger RNA by the addition of a cap structure is a key process that provides protection against nucleases and facilitates the export and translation of mRNA (1-4). The cap is formed on the first transcribed nucleotide of the pre-mRNA as a result of three consecutive enzymatic reactions: (i) the 5’ end triphosphate of pre-mRNA is hydrolysed to diphosphate by a 5’-triphosphatase; (ii) GMP is added by the RNA guanylyltransferase to create the cap intermediate GpppN and (iii) the RNA guanine-N7 methyltransferase (RNMT) catalyses the transfer of a methyl group from S-adenosylmethionine (AdoMet) to GpppN to create the mature cap, m7GpppN, and byproduct, AdoHcy (S-adenosyl homocysteine) (Figure 1A) (2). This capping mechanism targets the nascent RNA polymerase II transcripts and is conserved amongst eukaryotes and viruses, although structural organization of enzymes varies between organisms, e.g. the capping process can be carried out by a single complex or multiple enzymes (2, 5).

**Figure 1.**
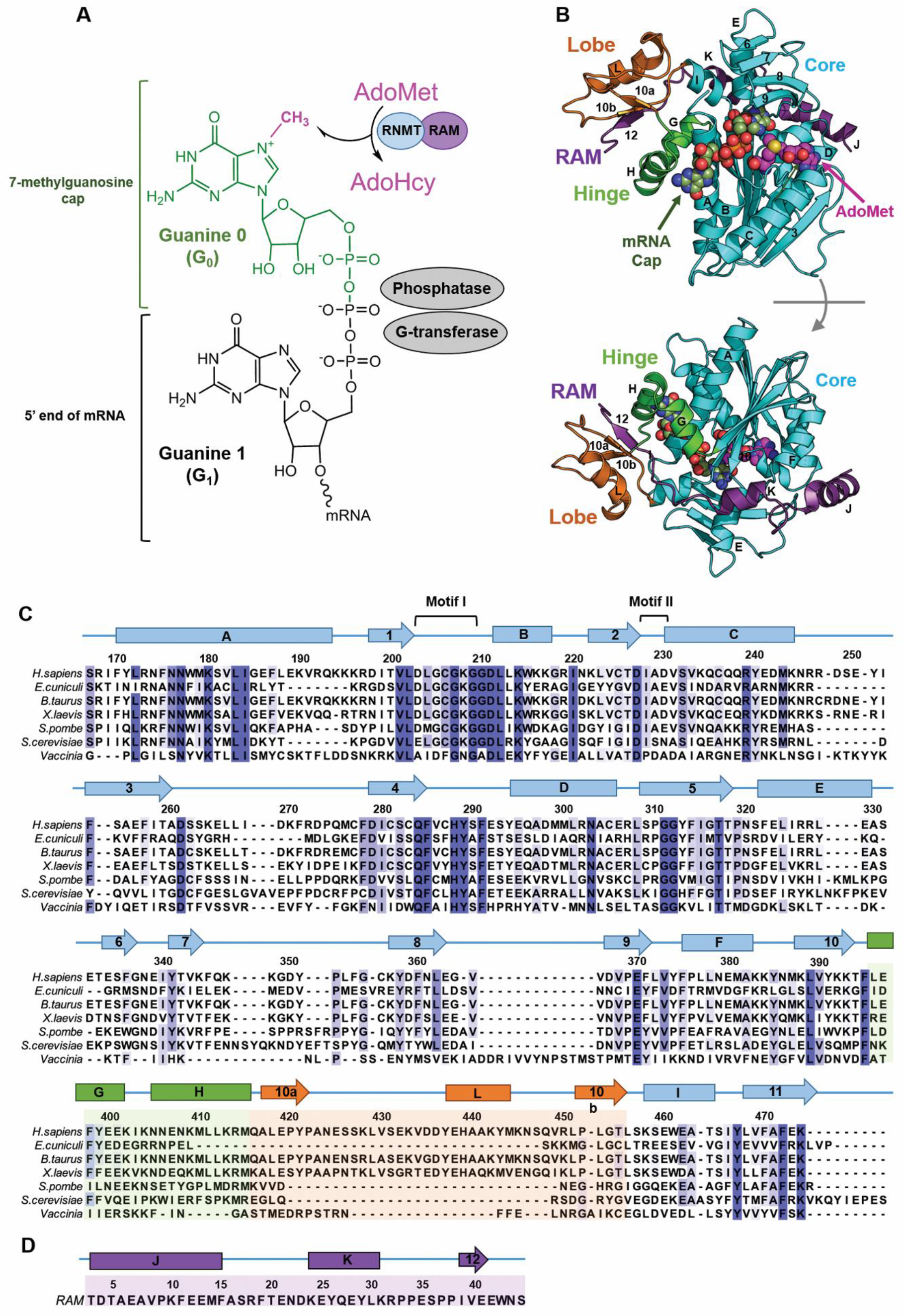
mRNA cap and mRNA methyltransferases. (A) Structure of mRNA cap and mRNA capping enzymes. The G_0_, G_1_, and methyl (CH3) groups of the cap are labelled. Three enzymes involved in a stepwise formation of the cap, namely G-transferase, phosphatase and methyltransferase (RNMT), are indicated next to the respective chemical group. RNMT catalyses transfer of a methyl group from AdoMet to G_0_ at N7 position, producing a by-product AdoHcy. (B) Structure of the human RNMT-RAM complex (PDB ID: 5E8J). RNMT core (residues 165–476) is shown in cyan, the α-helix hinge (residues 395-415) in green, and the modular lobe (residues 416-456) in orange. Nomenclature of key helices and strands is shown. RAM (residues 2–45) is displayed in violet. Representative binding poses of AdoMet and cap (G_0_pppG_1_) are shown using the spheres representation (also see Figure 3). (C) Multiple sequence alignment of human RNMT with its orthologs from a selection of eukaryotic organisms and viruses. The nomenclature of structural features shown was kept in line with Fabrega et al (12) and Varshney et al. (8). The elements of the RNMT core are shown in cyan, α-helix hinge - in green, and modular lobe - in orange, consistent with (B). (D) Sequence and secondary structure elements of RAM (residues 2-45).

In vertebrates, RNMT is activated by the RNMT-activating mini-protein (RAM) which has been shown to be required for efficient cap methylation, mRNA translation and cell viability (6). Grasso et al. demonstrated that regulation of RNMT-RAM complex has a crucial role in the gene expression changes needed during cell differentiation and reprogramming. In particular, the RNMT-RAM complex is important for pluripotency-associated gene expression. RAM is highly expressed in embryonic stem cells; during neural differentiation, RAM is repressed while high RNMT levels are maintained contributing to down regulation of pluripotency-associated genes and the emergence of the neural phenotype (7).

Recently, we reported the crystal structure of the catalytic domain (residues 165-476) of human RNMT in complex with the RNMT activation domain of RAM (residues 2–45) and AdoHcy (PDB ID: 5E8J) (8). Human RNMT consists of a catalytic domain (residues 121-476), homologous to other eukaryotic cap methyltransferases, and an N-terminal regulatory domain (residues 1-120) that facilitates the recruitment to RNA polymerase II transcription initiation sites (9, 10) and regulates RNMT activity (11). The presence of RAM permitted the stabilization and crystallization of a modular lobe of RNMT (residues 416–456), which was unresolved in previously reported structures of isolated RNMT (PDB IDs: 3BGV, 5E9W). This strongly suggested that the lobe is disordered in the absence of RAM.

The human RNMT has notable structural similarity to *E. cuniculi* mRNA cap methyltransferase Ecm1 despite having <40% sequence identity (12); the only major structural difference is the absence of the modular lobe in Ecm1. The core of the catalytic domain of RNMT displays the canonical Class I methyltransferase fold with alternating β strands (β1-β7) and α helices (αA-αE) (Figure 1B & C) (13). The lobe and an α-helix hinge containing helices αG and αH (residues 395-415) constitute a positively charged groove that accommodates the RNMT activating domain of RAM (residues 1-45) (8). In particular, the extended RAM region (residues 32-42) establishes several polar and hydrophobic interactions within the lobe and the α-helix hinge, while the more ordered N-terminal end of RAM (residues 1-31) interacts with helices αE, αD and β7-β8 loop.

Our recent findings showed that RAM was key to enhance the methyltransferase activity of RNMT despite its distal location to the active site of the enzyme (6, 8, 14). The results obtained from crystal structures, biochemical assays and molecular dynamics (MD) simulations suggested that RAM is critical for maintaining the stability and structure of the RNMT lobe. This stabilization promotes the binding of AdoMet and induces a significant increase in the methyltransferase activity in the presence of RAM as a result of a series of interactions affecting the α-helix hinge, α-helix A and multiple active site residues (8). However, details of the allosteric mechanism by which RAM activates RNMT and promotes recruitment of RNMT substrates - AdoMet and cap - remain elusive. Moreover, the available crystal structures of RNMT do not provide a clear insight into the binding mode of the substrates: the human RNMT-RAM complex and *Vaccinia* virus mRNA methyltransferase were solved only in the presence of the byproduct AdoHcy (8, 15). A set of crystal structures was reported for Ecm1 that included the methyltransferase in complex with individually bound ligands (AdoMet, AdoHcy and the cap analogue); however, the optimal poses of two substrates simultaneously bound could not be clearly revealed (12). Despite the concerted efforts of x-ray crystallography, biochemistry and computational biophysics in the attempt to understand RNMT activity and regulation, the binding modes of the RNMT substrates as well as the allosteric mechanism of RNMT activation by RAM remain unresolved issues. Therefore, an alternative approach needs to be taken in order to elucidate these key aspects of RNMT functioning. In this work, we shed light on these questions through a comprehensive computational study, employing a combination of several molecular simulation methods.

Over the past decade, MD simulations have become increasingly important in solving the atomic-level mechanisms of protein allostery, often revealing missing features that purely experimental techniques could not provide (16-19). However, a lack of sufficient sampling of events that take place on very large timescales constitutes a significant limitation when employing standard MD simulations to explore allosteric mechanisms. In order to overcome this problem, enhanced sampling simulations have shown to be extremely valuable for more efficient sampling of the conformational space of large bimolecular systems while using significantly less computational resources (20-25). In this study, we employ both standard (or *conventional*) molecular dynamics (cMD) and *accelerated* molecular dynamics (aMD) to: (i) elucidate the binding mode of RNMT substrates, cap and AdoMet, bound simultaneously to the enzyme’s active site and (ii) explain the allosteric mechanism by which RAM efficiently stimulates RNMT activity. Extensive simulations offer a detailed picture of the conformational changes occurring at the active site of the enzyme which promote binding of the reaction substrates and modulate the activity of RNMT. Furthermore, by performing community network analysis we provide evidence of the allosteric paths that connect RAM to the RNMT active site, thus offering the first comprehensive description of the molecular mechanisms of RNMT activation by RAM.

## MATERIALS AND METHODS

### Systems Setup

The crystal structure of the human RNMT in complex with RAM was recently reported by our group (8). The suite of crystal structures reported by Fabrega et al. elucidated the binding poses of AdoMet, AdoHcy and cap to *E. cuniculi* Ecm1 methyltransferase, although the simultaneous binding mode of both substrates remained elusive (12). Using these structures, we constructed a set of simulation systems to selectively investigate i) the binding modes of the substrates in the RNMT active site and ii) the allosteric activation of RNMT by RAM. A summary of systems and all molecular dynamics simulations performed in this work is provided in Table1.

**Table 1.** Summary of molecular dynamics simulations performed in this work.

| System | cMD | aMD |
| --- | --- | --- |
| apo RNMT | 100 ns | 200 ns |
| apo RNMT-RAM | 100 ns | 200 ns |
| apo RNMT-RAM (res 20-45) | 100 ns | 200 ns |
| RNMT-RAM · Cap + AdoMet | 100 ns | 200 ns |
| RNMT-RAM · Cap | 100 ns | 200 ns |
| RNMT · Cap + AdoMet | 100 ns | 200 ns |
| Total simulation time: | 600 ns | 1.2 μs |

#### RNMT-RAM complex with substrate(s)

Both substrates - AdoMet and the cap analogue G_0_pppG_1_ (‘cap’ throughout this paper) (Figure 1A) - were modelled and docked simultaneously into the human RNMT (residues 165-476) in complex with the full-length RAM (residues 2-45) using as references the crystal structures of RNMT-RAM with the bound AdoHcy (PDB ID: 5E8J) (8) and *E. cuniculi* mRNA cap methyltransferase Ecm1 solved with the cap (PDB ID: 1RI2) (12). Within the cap analogue, ‘G_0_’ corresponds to the guanine to be methylated by RNMT and ‘G_1_’ corresponds to the first nucleotide within the pre-RNA transcript. One of the simulated systems contained only the cap bound to RNMT-RAM.

#### apo-RNMT +/−RAM

Human RNMT in the absence of substrates was built and simulated with and without RAM.

#### apo-RNMT +RAM (20-45)

The role of the RAM N-terminal region in the RNMT activation was investigated by simulating RNMT in complex with a *truncated* form of RAM (residues 20 to 45), in which the N-terminal region was removed.

All models were prepared with the LEaP utility from the AMBER14 suite of programs (26) using the ff14SB force field (27). The N-terminus of RNMT was capped with an acetyl group (ACE), and the N- and C-termini of RAM were capped with ACE and a methylated amino group (NME), respectively.

The geometries of the ligands were refined with Gaussian03 (28) at the HF/6-31G* level. The optimized geometries were used to calculate the electrostatic potential-derived (ESP) charges using the RESP methodology (29), as implemented in the *antechamber* module in Amber14 (26). The force field parameters for AdoMet were generated with *antechamber* module, using the general AMBER force field (GAFF) (30).

Each simulation system was immersed in a cubic box of TIP3P water molecules that extended at least 15 Å away from the protein, and then was neutralized by adding the appropriate number of counterions (31). This was followed by steepest-descent energy minimization to remove steric clashes.

### Molecular dynamics simulation protocols

All MD simulations were performed using the pmemd.cuda module of AMBER14. Periodic boundary conditions were used. The cut-off distance for the non-bonded interactions was 10 Å. Electrostatic interactions were treated using the smooth particle mesh Ewald method (32). The SHAKE algorithm was applied to all bonds involving hydrogens, and an integration step of 2.0 fs was used throughout (33). Each system was heated to 300 K over a 50 ps interval using a weak coupling algorithm (34). A 200 ps equilibration was performed to allow the solvent to redistribute around the positionally restrained protein. The system was subsequently allowed to evolve unrestrained at constant temperature (300 K) and pressure (1 atm) using the weak-coupling algorithm (34). Atomic coordinates were saved every 20 ps for further analysis. Each system was initially simulated using *standard* molecular dynamics for 100 ns.

### Accelerated molecular dynamics (aMD)

The activation mechanism of RNMT involves protein-protein binding, domain conformational changes and long-distance correlated motions that often occur over very long timescales that are difficult to reach by cMD. To overcome this limitation, *accelerated* molecular dynamics introduces a positive boost energy to the potential energy function, effectively reducing the height of energetic barriers and enhancing conformational sampling (35-37). Contrary to other enhanced sampling protocols such as metadynamics (38, 39), aMD does not require predefined collective variables to initiate the enhanced sampling and has been successfully applied in several biomolecular systems to significantly improve the sampling of long-time scale event including allosteric mechanisms (40-43).

In aMD, a boost potential, *ΔV(r)*, is applied when the average potential energy of the system, *V(r)*, is below a previously defined reference potential energy *E_P_*. The modified potential, *V*\**(r)*, used for MD simulations, is then given by eq 1:

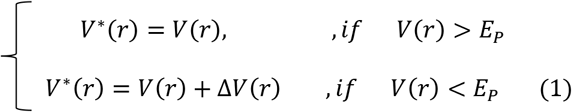

In this work, 200 ns aMD simulations were performed for each system (Table 1), using a “dual-boost” protocol. This included a boost potential energy applied to all the atoms in the system and an extra dihedral boost applied to the torsion angles:

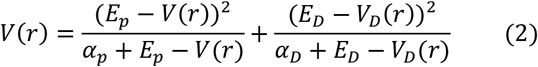

where *V_D_* is the dihedral energy, *E_P_* and *E_D_* are the reference potential and dihedral energies and *α_P_* and *α_D_* are the acceleration parameters that describe the strength of the boost for each term and determine the shape of the modified potentials. The parameters of the aMD simulation (*E_P_, α_P_, E_D_, α_D_*) were initially selected following the guidelines from previous works (43-45) and then further adjusted after preliminary aMD simulations (Supplementary Table S1).

### Analysis of MD trajectories

Protein structures and MD trajectories were visually inspected using the molecular visualization programs PyMOL (46) and VMD (47). Interatomic distances and angles, root-mean-square deviations and fluctuations (RMSD and RMSF) with respect to a reference structure and secondary structure analysis were monitored using the CPPTRAJ module in AmberTools 15 (26). Residue-per-residue cross-correlations analysis of the RNMT-RAM complex was performed using CPPTRAJ (26, 48) using the backbone Cα fluctuations. The two-dimensional schemes of the ligand-protein interactions were obtained using Ligplot+ (49). MDpocket (50) was used to evaluate the AdoMet and cap (G_0_) pocket volumes during the aMD simulation of each system. The two ligand pockets were defined in the reference system (RNMT-RAM with the bound cap and AdoMet) with MDpocket using 2000 snapshots from the aMD trajectory. The resulting grids were then superimposed on each simulated system and the volumes of the two ligands pockets were monitored along the 200 ns aMD trajectories. MDpocket was run using 10000 Monte Carlo iterations for each system. The minimum and maximum radii of the probe (“alpha”) spheres in each pocket were 3Å and 6Å, respectively. Electrostatic potentials were calculated using the Adaptive Poisson-Boltzmann Solver (APBS) with PyMol APBS tools (51).

### Unweighted Free Energy Landscapes (FEL)

Free energy landscapes (FEL) were constructed for each system to characterize the “open” / “closed” RNMT active site conformations sampled during the aMD simulations. Two collective variables (CVs) that could discriminate between several active site conformations were identified: CV1 describes the accessibility to the cap binding pocket and was defined as the centre of mass (COM) distance between the upper half of RNMT α-helix A (residues 2-11) and the β8-β9 loop (residues 362-366); CV2 describes the accessibility to the AdoMet binding pocket and was defined as the COM distance between Motif I (residues 205-209) and the β4-αD loop (residues 285-289), as shown in Figure 5A. The cap guanine pocket was considered to be accessible (“cap-open” state) when CV1 is >10 Å, which favoured binding of G_0_ to the cavity created between α-helix A and the β lid. The AdoMet binding pocket was considered to be open and accessible to AdoMet when CV2 was >11 Å. The “fully open” conformation of RNMT is only obtained when CV1 and CV2 are greater than 10 Å and 11 Å respectively, allowing the access of both substrates. A “closed” conformation is obtained when CV1 and CV2 are bellow 9Å and 10 Å respectively.

In principle, aMD simulations can recover the equilibrium properties of the unbiased potential by reweighting statistics using the Boltzmann factor of the boost potential e^(βΔV(r))^ (37). However, in aMD simulations of complex biomolecules, large values of boost potentials are normally used to gain sufficient sampling, which leads to a large “energetic noise” during the reweighting process (52, 53). Hence, unweighted FEL are presented in this work, which are employed only for comparison goals in line with previous aMD studies of large biomolecular systems (41, 54).

### Community Network Analysis

Allosteric networks in the RNMT-RAM complex were studied by means of community network analysis using the NetworkView plugin in VMD (55, 56). The protein system in the molecular dynamics simulations is represented as a set of “nodes” corresponding to each protein residue. Edges connecting pairs of nodes are drawn when two residues are within 4.5 Å of each other for at least 75% of the aMD trajectory, yielding a complex all-residue network. To obtain a simplified and easier to interpret coarse-grained representation, the resulting network map is decomposed into “communities” using the Girvan-Newman algorithm (57): large groups of strongly connected nodes (i.e. residues) are clustered into communities, which are then loosely connected to one another by edges. The edges are weighted to reflect strength (so-called cumulative betweenness) of the interactions between communities and thus offer a measure of potential allosteric interactions and pathways (58). The community networks are represented in diagrams displaying the communities as circles, with radii proportional to the number of residues comprising a community, connected by edges, whose widths are proportional to the strength of the intercommunity interaction.

## RESULTS

### Binding modes of AdoMet and cap to RNMT

Our first goal was to elucidate the reactive configuration of substrates in the active site of RNMT. Since the crystal structure of human RNMT (or homologous mRNA methyltransferases) in complex with two simultaneously bound substrates is not available, we have built a model by superimposing individual ligands from the Ecm1 methyltransferase set of structures onto the active site of human RNMT-RAM (see details in Methods). The obtained model of RNMT-RAM in complex with AdoMet and cap was then validated through a combination of cMD and aMD simulations. The complex was first relaxed for 100 ns using unrestrained cMD (Figure 2A).

**Figure 2.**
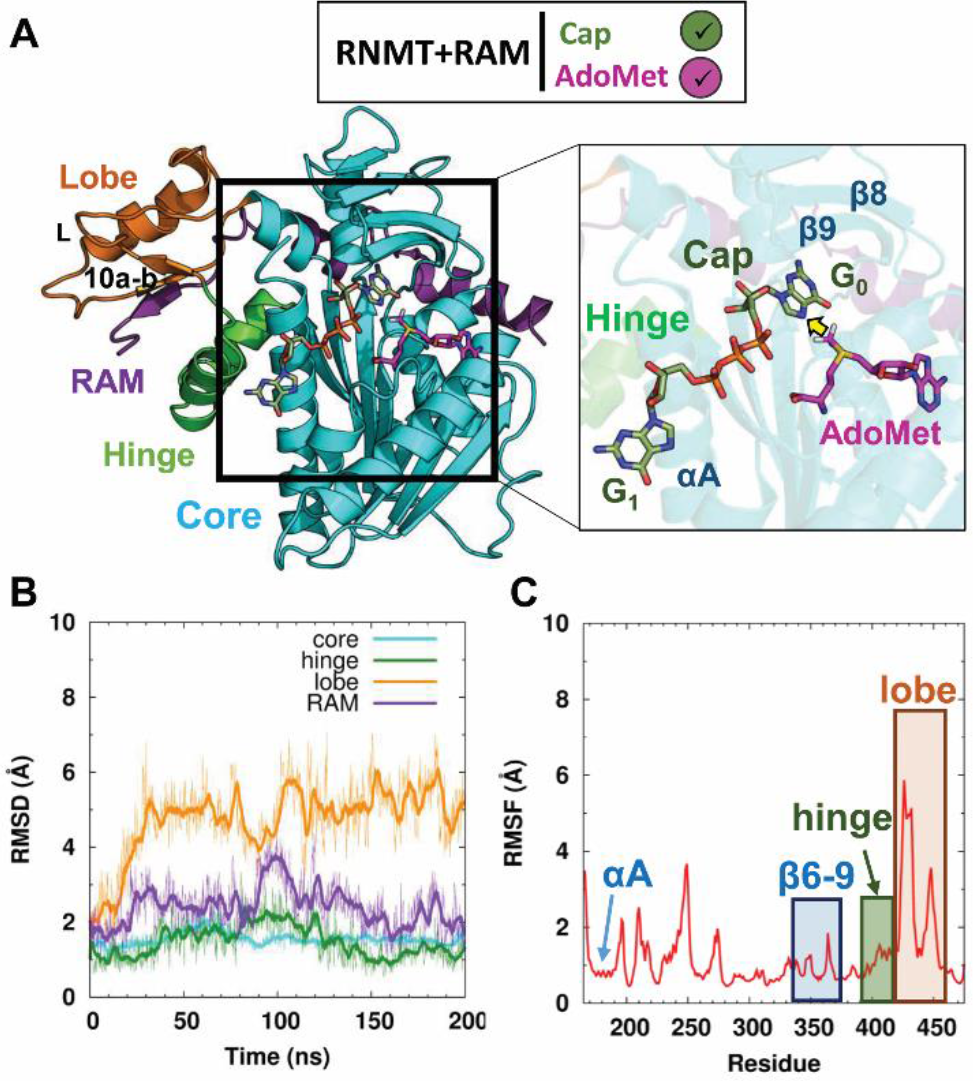
Characterisation of the RNMT-RAM complex in a simulation with bound AdoMet and cap. (A) Structural model of RNMT-RAM (cartoon representation) in complex with AdoMet and cap (stick representations, with magenta and green carbon atoms, respectively). A close-up view shows the ligands within RNMT in a methyl-transfer reactive conformation; the transferred methyl group is indicated by a yellow arrow. Relevant structural elements of RNMT are labelled. (B) Time evolution of backbone RMSD (in Å) of the key structural elements of RNMT during the aMD simulations. (C) RMSF values (in Å) of the RNMT residues obtained from aMD simulations.

In order to sample more efficiently the protein free energy landscape and possible allosteric effects, which often take place on long timescales, the relaxed system was simulated for 200 ns using aMD.

#### RNMT-RAM structure and conformational dynamics in the presence of ligands

aMD simulations of the RNMT-RAM complex in the presence of AdoMet and cap showed a high degree of stability of the key RNMT structural elements (Figure 2B). The core domain of RNMT (residues 165-476) remains stable throughout the trajectory (average RMSD 1.5 Å) and maintains structural features observed in the crystal structure of human RNMT-RAM (8). In particular, the α-helix A (residues 170-194), that is positioned adjacent to the cap binding site, keeps its original conformation (RMSF 0.8 Å), in which it interacts with the α-helix hinge element (residues 395-415) (Figure 2A and 2C). Strands β8 and β9 (residues 356-362, 365-371), which form a “lid” for the cap binding pocket similar to the one observed in the Ecm1 methyltransferase (12), also remain highly stable during the trajectory (RMSF 1.3 Å). The region displaying the highest degree of flexibility was the lobe (residues 416-456). That was mainly due to a long loop connecting strand β10a and α-helix L. Nonetheless, the secondary structure elements were maintained along the aMD trajectory in all regions of RNMT, including the lobe. The ordering of the lobe was a result of multiple polar interactions with the activation domain of RAM (residues 2-45), that binds to a positively charged surface groove, as detailed in our previous work (8). Thus, RAM introduces a high degree of stability of the catalytic domain of RNMT. Overall, the protein structure and conformational dynamics observed in the aMD simulation in the presence of AdoMet and cap were very similar to that of *apo*-RNMT-RAM in the crystal structure and our previous cMD study (8). Therefore, the obtained structural model, which incorporates simultaneously bound reaction substrates (AdoMet and cap), can be used to determine with atomic resolution their binding poses and interactions within the RNMT active site – an important information missing in the crystallographic structures available to date.

#### AdoMet binding mode

AdoMet binds into a pocket formed by strands β1 and β2, adopting a configuration common to AdoMet-dependent methyltransferase enzymes (8, 12, 13, 15). No significant differences were observed between the AdoMet conformation obtained in this study (Figure 3A; see also Supplementary Table S2) and the AdoHcy binding pose in the RNMT crystal structure (8).

**Figure 3.**
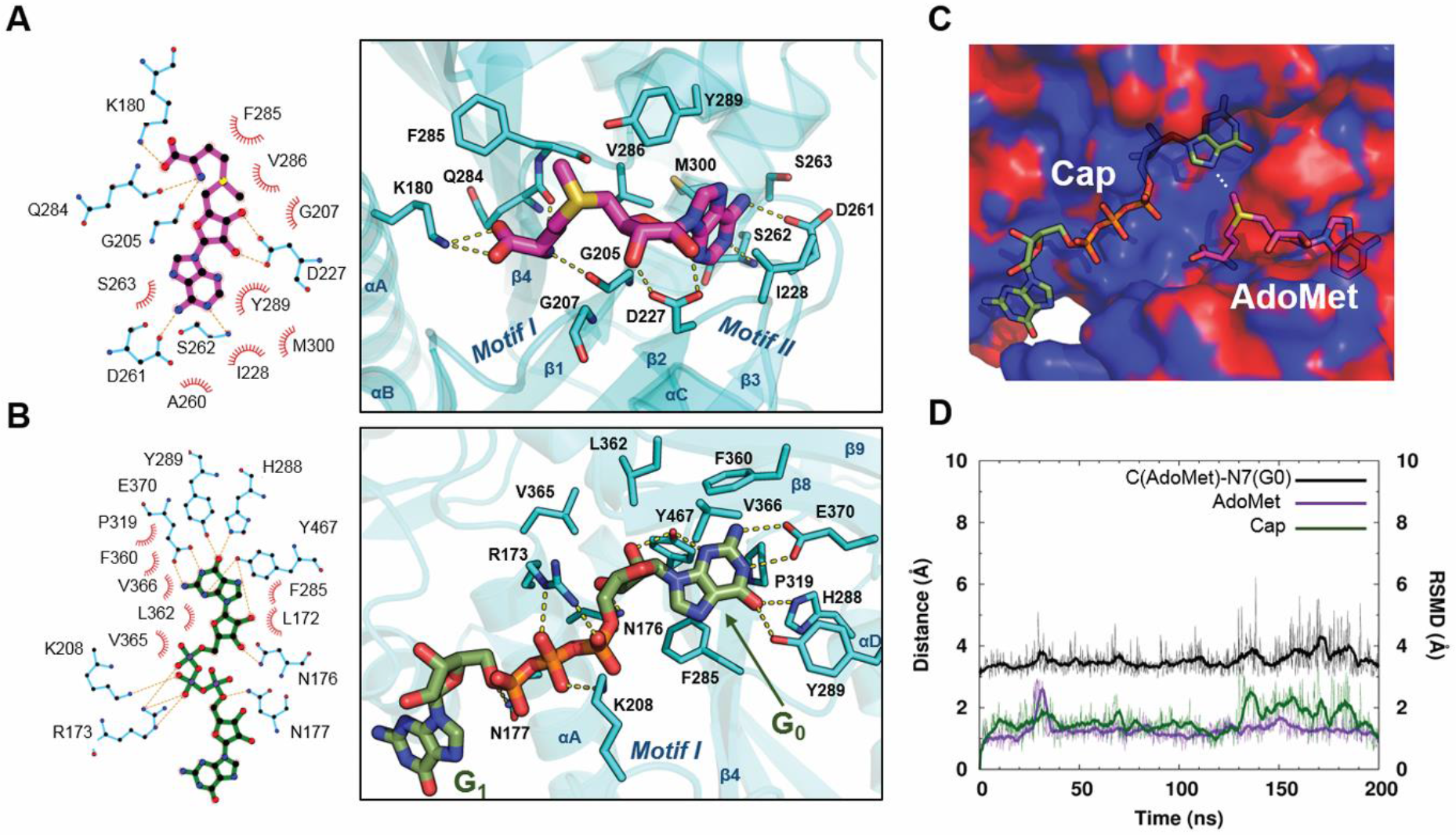
Binding modes of AdoMet and cap in the RNMT active site. (A) 2D protein-ligand interaction map of RNMT and AdoMet (magenta) (left), and a representative aMD snapshot of AdoMet bound to the RNMT active site (right). (B) 2D protein-ligand interaction map of RNMT and cap (green) (left), and a representative aMD snapshot of cap bound to the RNMT active site (right). (C) Electrostatic potential surface representation of the AdoMet and cap binding pockets. Isovalue surfaces of the electrostatic potential around the active site at values of −1 and +1 k_B_T e^−1^ are coloured red and blue, respectively. (D) Time-evolution of the distances between AdoMet (carbon atom of the methyl group) and the cap G_0_ (N7 atom) (black line) in the aMD simulation; the monitored distance between the groups is indicated by a white dotted line in panel (C). The RMSDs of AdoMet (magenta) and cap (green) are plotted against the secondary (right) vertical axis.

The carboxylate group establishes a salt bridge interaction with K180, a known key residue for RNMT activity (8),which is located within the α-helix A. The amino acid moiety of AdoMet also interacts with the glycine-rich loop that connects strand β1 and α-helix B and includes the conserved “Motif I” (Glu/Asp-X-Gly-X-Gly-X-Gly; residues 203-210) (Figure 1C). The amino group of AdoMet forms a hydrogen bond with the carbonyl oxygen of G205. In addition, the 2’ and 3’-hydroxyls of the adenosine ribose interact with D227 within the “Motif II” loop (residues 227-230), and non-polar contacts are established with I228. The long loop connecting strands β3 and β4 has an important role in stabilizing AdoMet by establishing two hydrogen bonds between the adenine atoms N6 and the side chain of D261 as well as between the adenine N1 and the backbone N of S262. Q284 in strand β4, establishes a hydrogen bond with the amino group of AdoMet whereas V286 within the β4-helix αD loop stabilizes the adenine base through hydrophobic interactions with V286. As a result, AdoMet is stabilized within a pocket displaying a negative electrostatic potential due the presence of D227 and the carbonyl oxygens of Q284 and F285, which is also favourable for the stability of the AdoMet sulfonium ion (Figure 3C). It is important to note that the above-mentioned residues involved in binding of AdoMet are highly conserved (Figure 1C), highlighting that the catalytic domain of RNMT has high sequence similarity and shares structural and functional features with similar mRNA methyltransferases.

#### Cap binding mode

G_0_ of the cap binds within a pocket located between the β4 strand-α-helix D loop, which constitutes the “floor” of the pocket, and strands β8-9 that form a lid over the cap binding site (Figure 3B), in a similar fashion to what was suggested for Ecm1 (12). The active site residues establish a number of non-polar and hydrogen-bonding and salt-bridge interactions with the cap groups (Supplementary Table S2). F285 establishes a stacking interaction with G_0_. On the right-hand side of the active site, H288 (Nε) and Y289 (OH) form two hydrogen bonds with the guanine O6. P319 contributes to guanine stabilization by establishing hydrophobic interactions with the cap. Key hydrophobic interactions are observed between G_0_ and β-lid residues F360, L362, V365 and V366. The end of the lid is formed by E370 that establishes two hydrogen-bond interactions with the G_0_ N1 and N2. Importantly, Y467 located at the back of the pocket, establishes two hydrogen bonds with N3 and O2’. The left-hand side of the pocket is formed by key residues located on the α-helix A: R173 forms several polar interactions with the polyphosphate chain connecting G_0_ with G_1_, whereas Nδ’s of N176 and N177 form hydrogen bonds with the ribose O3’ and the cap phosphate groups, respectively. Interestingly, K208, located in the key loop responsible for AdoMet binding (“Motif 1”), establishes two salt bridges with α and β phosphates of the mRNA cap. Finally, the electrostatic surface representation of the cap binding pocket revealed an extended, highly positive region fitting the polyphosphate chain of the cap, with contributions from R173, K180, K208, R239 and R246 (Figure 3C). The G_0_ pocket displays a more neutral surface which is favourable for substrate binding. Similar to the AdoMet pocket, the residues involved in binding of the G_0_ and polyphosphate chain of the Cap are highly conserved (Figure 1C).

The above-detailed multiple RNMT-ligand interactions enabled tight binding of AdoMet and cap in the presence of RAM, as demonstrated by a very high degree of stabilization of both ligands during long *unrestrained* cMD and aMD simulations: the average RMSD values were about 1.5 Å for both ligands (Figure 3D); ligand dissociation or at least substantial deviations from the original structure were not observed. In addition, the ligands maintain the conformation optimal for the AdoMet-to-cap methyl transfer (Figure 3D): the distance between the AdoMet CH3 and the methyl acceptor (G_0_ N7) was maintained at around 3.7 Å (a structure of the representative snapshot is included in Supplementary Data in the PDB format). During the cMD and aMD simulations, no direct contacts were observed between RNMT and the N7 atom of G_0_, the sulphur atom of AdoMet or the methyl group of AdoMet, in line with the models suggested by Fabrega et al. (12).

In order to explore possible origins of the allosteric effect of RAM on the substrates binding, an additional system, namely RNMT with the bound AdoMet and cap but in the absence of RAM, was simulated using the same protocol; the results are shown in Supplementary Figure S1. In terms of ligands interactions and dynamics, this system displayed very similar behaviour to that in the presence of RAM (Figure 3D), showing identical ligand binding poses and interactions with the active site residues, as well as similar RMSD values for both ligands; the methyl-transfer distance was also maintained. This illustrates that RAM does *not* affect stability of ligands in the active site once they are *bound* there. It should be mentioned that this does not rule out a direct role of RAM in the subsequent catalytic mechanism, e.g. through electrostatic interactions (59). In the RNMT structure, as expected, the only major change was the disordering of the modular lobe region which occurs in the absence of RAM, whereas α-helix A, the α-helix hinge and strands β6-9 maintained the same overall conformations.

In summary, cMD and aMD simulations offered a detailed picture of the AdoMet and cap binding poses and interactions within the RNMT active site, which complements and is consistent with previous structural and biochemical data. This provides the most complete description of the binding modes of the substrates within the RNMT active site to date. Nonetheless, and despite the stabilization effect observed by RAM on the overall RNMT structure and dynamics, the mechanism by which RAM regulates RNMT activity remained unclear. Hence, we next investigated whether RAM can allosterically regulate RNMT activity through selection of specific RNMT conformations that could enhance binding of AdoMet and cap to the active site.

### RAM stabilizes RNMT and enhances the active site conformations promoting the substrates binding

As our simulations revealed no effect of RAM on the poses and interactions of two substrates when *bound* in the RNMT active site, next we explored an alternative explanation of the RNMT activity modulation by RAM: we investigated whether the role of RAM is to *prepare* the AdoMet and cap pockets for optimal binding of the ligands. We carried out cMD and aMD simulations of *apo*-RNMT in the presence and absence of RAM, focussing on the conformational changes in the previously characterised AdoMet and cap binding pockets. Furthermore, recent experimental data suggested that the cap binding could promote the binding of AdoMet to the RNMT-RAM complex following a co-operative binding model (8). To provide a molecular rationale for this observation, RNMT-RAM was also simulated in the presence of the cap only.

#### RAM modulates overall stability of RNMT

Analysis of *apo*-RNMT dynamics provided valuable insights into the allosteric effects induced by RAM (Figure 4).

**Figure 4.**
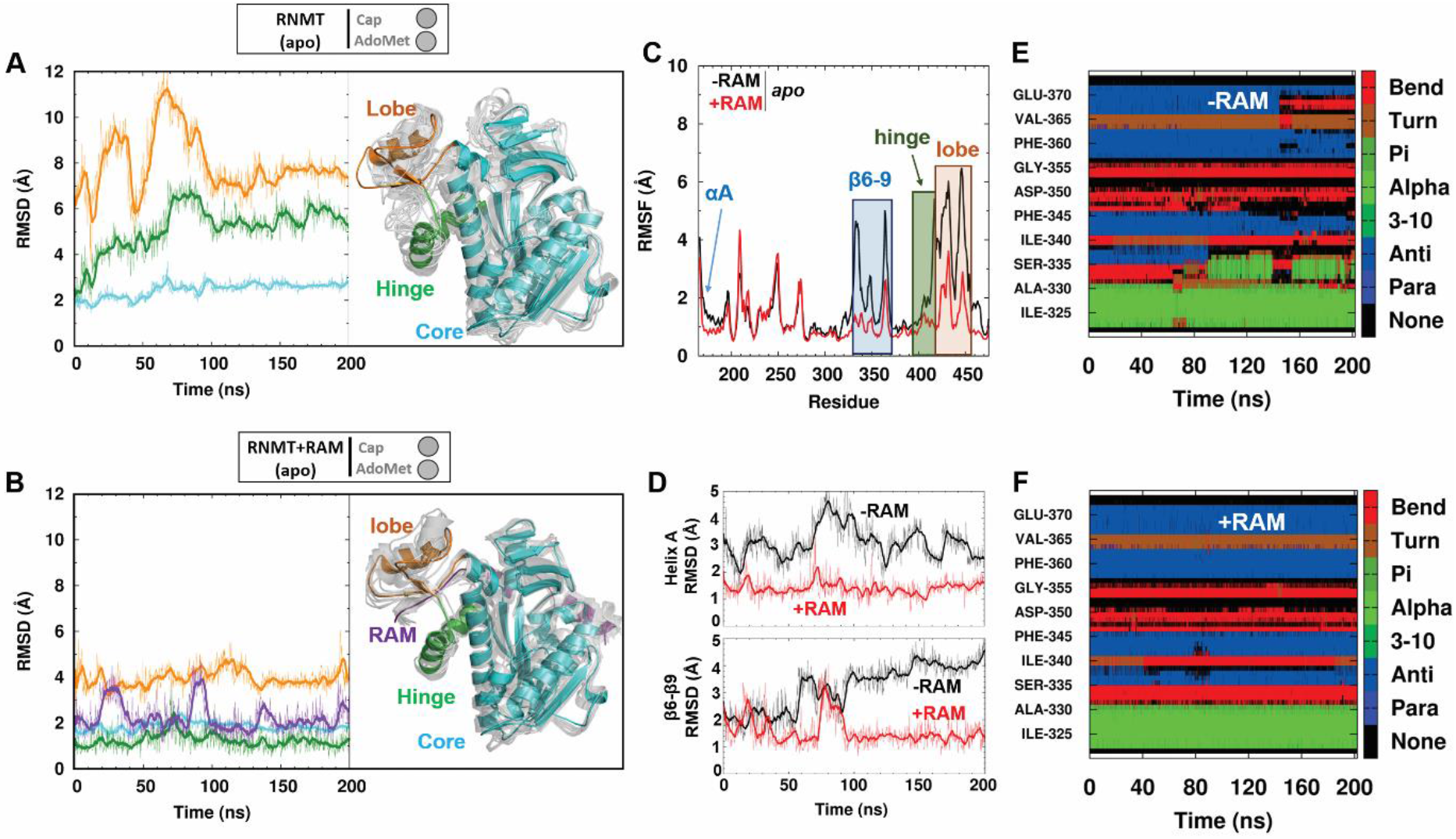
Modulation of RNMT stability by RAM. (A-B) Left: Time evolution of the backbone atoms RMSD (in Å) of key structural elements of RNMT during the aMD simulation of *apo*-RNMT in the absence (A) and presence (B) of RAM. Right: A representative snapshot of *apo*-RNMT in the absence (A) and presence (B) of RAM, displaying the key structural elements: catalytic core (cyan), hinge (green) and lobe (orange) regions. The representative snapshot is superimposed with multiple structures (light grey) that were sampled from the entire aMD trajectory and illustrate the range of conformational dynamics in each system. (C) The backbone RMSFs (in Å) of the catalytic domain of RNMT (residues 165-475) computed from the aMD simulation of *apo*-RNMT in the absence (black) and presence (red) of RAM. (D) Time evolution of the backbone RMSD (in Å) of α-helix A and strands β6-9 during the aMD simulation of *apo*-RNMT in the absence (black) and presence (red) of RAM. (E-F) Secondary structure analysis of the β-strand ‘lid’ region of RNMT (residues 320-375) performed on the aMD trajectories of *apo*-RNMT in the absence (E) and presence (F) of RAM.

The RMSFs and RMSDs were computed from the aMD simulations of RNMT in the presence or absence of RAM. Consistent with the crystal structure and our previous cMD simulations of RNMT-RAM (8), the lobe region displays the highest flexibility in either system, though it is significantly higher in the absence of RAM (average RMSD of the lobe backbone atoms were 4.5 Å vs 8.0 Å with and without RAM, respectively) (Figures 4 A-B). The calculated backbone RMSF values also highlight the stability brought about by RAM within the lobe region (Figure 4C). This disorder-to-order transition is achieved as a result of multiple RNMT-RAM interactions that enabled the crystallization of the modular lobe of RNMT only in the presence of RAM (8). The degree of flexibility of the lobe is directly correlated with that of the α-helix hinge, which remained remarkably stable in the RNMT-RAM simulation (Figure 4C). Conversely, disordering of the lobe in a simulation of RNMT triggered by the absence of RAM, notably increased flexibility of the hinge and the upper half of α-helix A, which contains the important active site residues N176 and K180 (Figure 4D). Flexibility of strands β6-β9, which form the active site “lid”, was also significantly pronounced in the absence of RAM. Secondary structure analysis showed a progressive loss of strand elements in the β6-β9 region during the aMD simulation of RNMT, while they were maintained in the simulation of the RNMT-RAM complex (compare Figures 4E and 4F). Large flexibility of β8-β9 loop also triggered disordering of strand β9, located in the upper region of the cap binding pocket and containing the residues V365, V366 and E370, which are important for binding of cap.

#### RAM selects the RNMT active site conformations optimal for substrates binding

Next, we investigated whether the observed RAM-modulated stabilization of various structural elements in *apo*-RNMT is also linked to changes in the active site, e.g. through promotion of certain conformations that favour binding of substrates. In order to quantify such allosteric effect, we introduced two descriptors of the substrate binding pockets, namely collective variables (CVs) 1 and 2 (Figure 5A).

**Figure 5.**
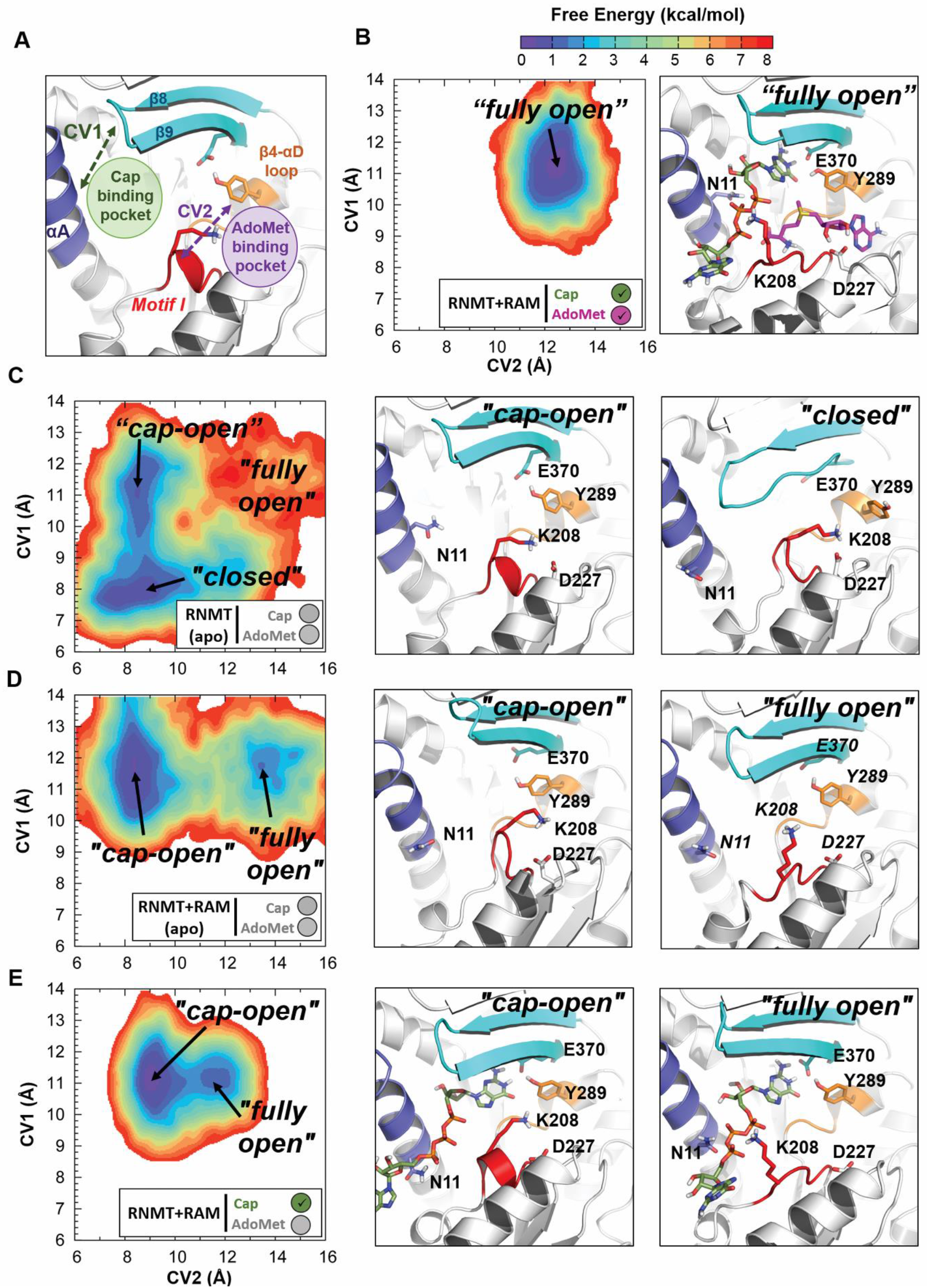
RAM-dependent modulation of the RNMT active site conformations. (A) Definition of collective variables (CVs) used to construct the free energy landscape (FEL) of RNMT. CV1 describes the cap binding pocket accessibility and is defined as the centre of mass (COM) distance between the upper half of the RNMT α-helix A (residues 2-11, highlighted in dark-blue) and the β8-β9 loop (residues 362-366, in cyan). CV2 describes the AdoMet binding pocket accessibility and is defined as the COM distance between Motif I (residues 205-209, in red) and the β4-αD loop (residues 285-289, in orange). (B-E) The conformational FELs of the active site of RNMT calculated for the simulation systems: (B) RNMT-RAM complex with the bound substrates AdoMet and cap; (C) *apo*-RNMT; (D) *apo*-RNMT in complex with RAM; and (E) RNMT-RAM complex with the bound cap. The FELs were built using all the snapshots from the corresponding aMD simulation and are shown as 2D contour plots with the lines in 0.5 kcal/mol increments. The colour scale corresponding to (unweighted) free energies is shown at the top. The simulation system is indicated by a legend in the box below the FEL. The most sampled conformational states, corresponding to the FEL minima, are labelled, and the corresponding representative structures of the RNMT active site are shown in the middle and right panels, next to the FELs.

The two CVs were used to build free energy landscapes (FEL), which in a simple manner represent all the conformations of the active site of RNMT that were sampled during an aMD simulation and provide a straightforward characterisation of the RNMT active site.

As a reference we used the structural model of RNMT-RAM with the bound AdoMet and cap ligands, which naturally provides the optimal conformations of each substrate pocket and of the entire active site (“*fully open*” conformation) (Figure 5B). The FEL obtained for this system shows a single free energy minimum, i.e. the conformation that was predominantly sampled during the simulation, which is located at CV1=11.5 Å; CV2=12.2 Å - a “*fully open*” configuration favouring the binding of both substrates. In this conformation, the stability displayed by α-helix A and β8-β9 strands enables the correct docking of G_0_. Motif I is stabilized by the K208-cap phosphate interaction, whereas Y289 interacts with G_0_, E370 and the adenosine base of AdoMet. D227 in Motif II also interacts with the ribose hydroxyls of AdoMet. Overall, both AdoMet and cap binding pockets were strongly stabilized and sampled only in the “*fully open*” conformation, as expected for the simulation in the presence of substrates.

The FEL of *apo*-RNMT in the absence of RAM displayed a global minimum located at CV1=7.8Å, CV2=9Å (“*closed*” conformation) (Figure 5C). In this conformation, the cap pocket is collapsed as a result of the displacement of α-helix A towards the guanine binding area and the partial loss of some secondary structure in the β lid (strands β8 and β9). The disordering and increased flexibility of strand β9 affects the position of the sidechain of Y289 that then flips towards the AdoMet binding site. In addition, the sidechain of K208 reorients towards the electronegative AdoMet pocket and interacts with D227. As a result, the Motif I loop adopts an alternative conformation that obstructs AdoMet accessibility to the active site, thus leading to the “*closed*’ conformation. A second local minimum in this FEL is located at CV1=11 Å, CV2=8.5 Å, representing the “*cap-open*” conformation (Figure 5C). This configuration would allow binding of cap as a result of a more favourable position of α-helix A and preserved structure of strands β8-β9. In contrast, accessibility for AdoMet to its pocket is limited: similar to the “*closed*’ conformation, Motif I loop is oriented towards the AdoMet binding site, blocking the access to AdoMet, despite the fact that Y289 is stabilized by E370. Altogether, this results in a conformation of the RNMT active site that offers binding of cap but not AdoMet, thus the “*cap-open*” state. The population of this state is slightly lower than of the “closed” state, and the barrier on the path between the two is low (1.5 kcal/mol). Thus, the FEL illustrates a dynamic equilibrium between these two states as sampled in the aMD simulation. The “*fully-open*” conformation was poorly sampled in the aMD simulation of *apo*-RNMT (Figure 5C) suggesting that in the absence of RAM simultaneous binding of AdoMet and the cap is strongly disfavoured.

The FEL obtained for the *apo*-RNMT in the presence of RAM showed a strikingly different picture (Figure 5D): in addition to the “*cap-open*” state (located at CV1=11.9 Å, CV2=8.5 Å), which now represents a global minimum (i.e. the most populated conformation), the “*fully open*” state (CV1=11.9 Å, CV2=13.5 Å) was now well sampled, while the “closed” state completely disappeared. In the “*cap-open*” conformation firmness of the cap pocket is facilitated by the Y289 interaction with E370, which stabilizes strand β9 of the β-lid. Similar to the *apo*-RNMT (no RAM) system, K208 is still flipped towards D227, inducing a conformational change in Motif I loop which limits accessibility to the AdoMet pocket in the “*cap-open*” state. In contrast, in the “*fully-open*” state, the Motif I loop shifts towards the cap binding site, increasing the accessibility to the AdoMet pocket; the cap pocket still offers conformation suitable for the cap binding. The “*fully open*” state displays a broader and more shallow landscape compared to the “*fully-open*” minimum in the presence of substrates (compare Figures 5D and 5B). This confirms that presence of the substrates stabilizes and significantly reduces the flexibility of the pockets. Moreover, the FEL showed a notable flattening of the region separating the “*cap-open*” and “*fully-open*” conformations compared to the *apo*-RNMT system in the absence of RAM (Figure 5D *vs* 5B). This indicates that transitions from the “*cap-open*” to the “*fully-open*” states are kinetically allowed in *apo*-RNMT-RAM.

Our data reveal that in the presence of RAM, *apo*-RNMT will be predominantly found in a state suitable for cap binding (the “*cap-open*” state), while in the absence of RAM the “*cap-open*” state first needs to be reached from the “*closed*’ state. Importantly, the “AdoMet-open” conformation (with the cap pocket blocked) was never sampled in any of the studied systems. Altogether, our results show that RAM allosterically primes the RNMT active site for the cap binding. Furthermore, this strongly suggests the likely scenario in which the cap binding occurs first, and this promotes the subsequent AdoMet binding. This hypothesis is in agreement with our previous fluorescence polarisation experiments that suggested that binding of the cap can enhance binding of AdoMet (8).

Additional support to this finding is provided by the results for the RNMT-RAM-cap system, which was simulated with the bound cap (but no AdoMet) using the same protocol (Figure 5E). As expected, the FEL showed a global energy minimum in the “*cap-open*” state (CV1=11 Å, CV2=9 Å), which naturally favours the cap binding. The “*fully open*” conformation was also sampled although it can be seen (Figure 5E) that conformational space in this case was more restricted as a result of the extra stabilization of the active site introduced by the presence of the cap. In this state K208 interacts with the polyphosphate chain of the cap, while the presence of the bound guanine (G_0_) greatly stabilizes Y289. Both interactions ensure access to the AdoMet pocket, and thus the conformation favours the binding of AdoMet. Importantly, substantial flattening of the entire landscape as well as of the barrier between the “cap-open” and “fully-open” states was observed and confirms the hypothesis that binding of the cap facilitates binding of AdoMet.

The distribution of the volume of the substrate pockets during the simulations was used as an additional quantitative measure to provide further details about the substrate binding mechanisms (Figure 6).

**Figure 6.**
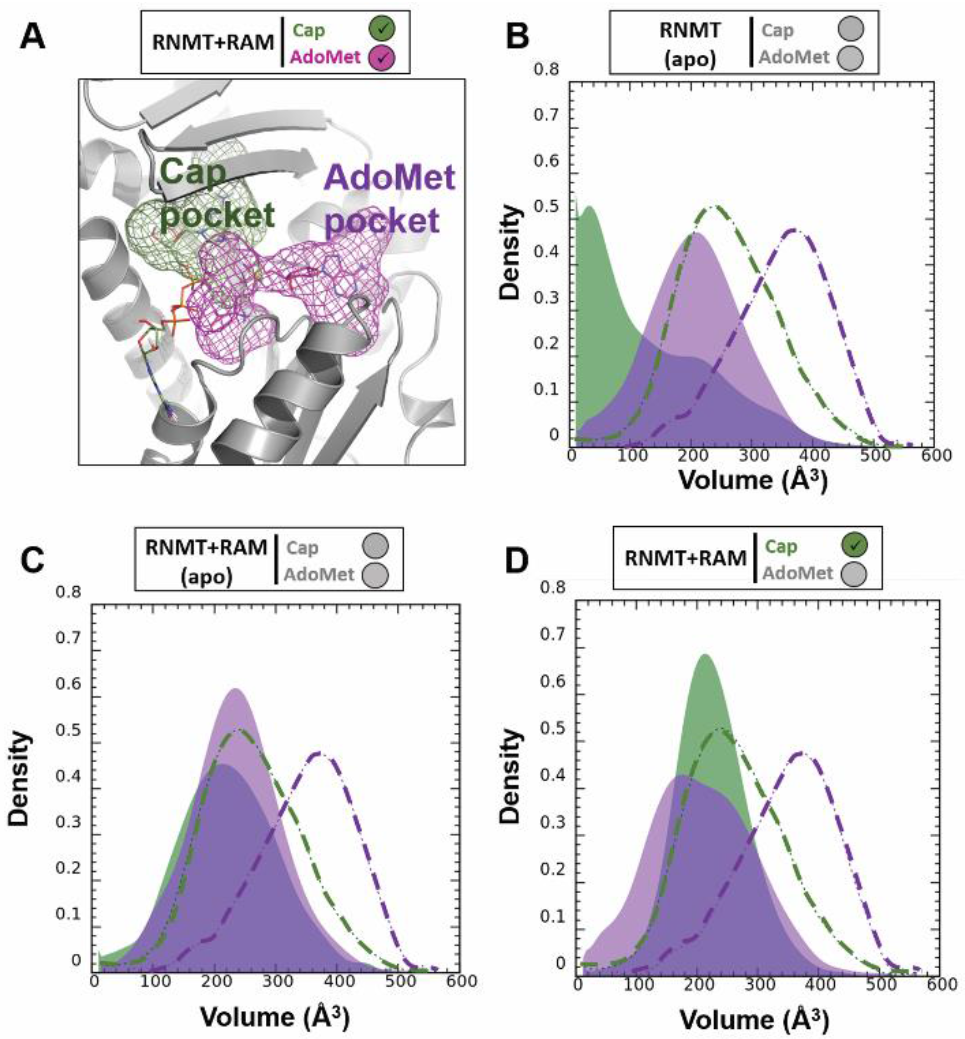
Distribution of the volume of the cap and AdoMet binding pockets. (A) The binding pockets of the cap (green) and AdoMet (magenta) were defined for the reference system (RNMT-RAM in the presence of both substrates) using MDpocket (see details in the Methods) and are shown as mesh surfaces. The ligands are shown as sticks inside the mesh surfaces. (B-D) Distribution of the volume of the cap (green) and AdoMet (magenta) binding pockets during the aMD simulations of: (B) *apo*-RNMT; (C) *apo*-RNMT in complex with RAM; and (D) RNMT-RAM with the bound cap. The distributions obtained from the simulation of the reference system (RNMT-RAM with both substrates) are shown as dashed lines on each plot for comparison.

This complements the free energy landscapes and conformational states reported in Figure 5. In *apo*-RNMT, the volumes of the both cap and AdoMet pockets are significantly reduced (thus collapsed, inaccessible pockets) when compared to those from the reference system (RNMT-RAM with the bound substrates) (Figure 6B). The presence of RAM significantly increases the volume of the cap pocket, whereas the AdoMet binding site remained largely inaccessible (Figure 6C). As expected, the binding of the cap ensured the optimal volume values of the G_0_ pocket (Figure 6D). Interestingly, in the presence of the cap, the volume of the AdoMet binding pocket was still significantly lower than in the reference system. This is consistent with the fact that the preferred conformation in the cap-bound system was the “cap-open” conformation and not the “*fully-open*” state (see Figure 5E). We can also interpret this as an indication of a plausible induced-fit binding mechanism for AdoMet, which is triggered by the K208 reorientation towards the cap polyphosphate chain.

Overall, analyses of the free energy landscapes and volumes of the binding pockets reveal that presence of RAM significantly enhances population of the “*cap-open*” and “*open*” conformations of RNMT, which are suitable for binding of substrates and particularly favourable for binding of the cap. This is consistent with the conformational selection mechanism of allostery. Furthermore, binding of the cap to the RNMT-RAM complex favours an active site conformation that is suitable for AdoMet binding, thus providing a rationale to the cooperative binding model suggested previously (8). In striking contrast, in the absence of RAM, RNMT displays significantly higher conformational flexibility and the “*closed*’ conformational state of the active site is predominantly sampled, in which binding of substrates is hindered.

### An allosteric network mediates RNMT structural modulation by RAM

The results obtained from the aMD simulations suggest that RAM modulates the conformational changes in RNMT to alter its active site conformation. However, the precise mechanism of how this is achieved, i.e. specific interactions between RNMT and RAM and the way they are translated into conformational changes in the active site, remained unclear. In order to reveal the molecular origin of this regulatory effect and identify putative allosteric pathways, advanced post-processing analyses were performed on the aMD trajectories of the *apo*-RNMT-RAM complex; this included a residue-by-residue cross-correlation analysis, protein dynamic networks and a coarse-grained community network analysis (Figure 7).

**Figure 7.**
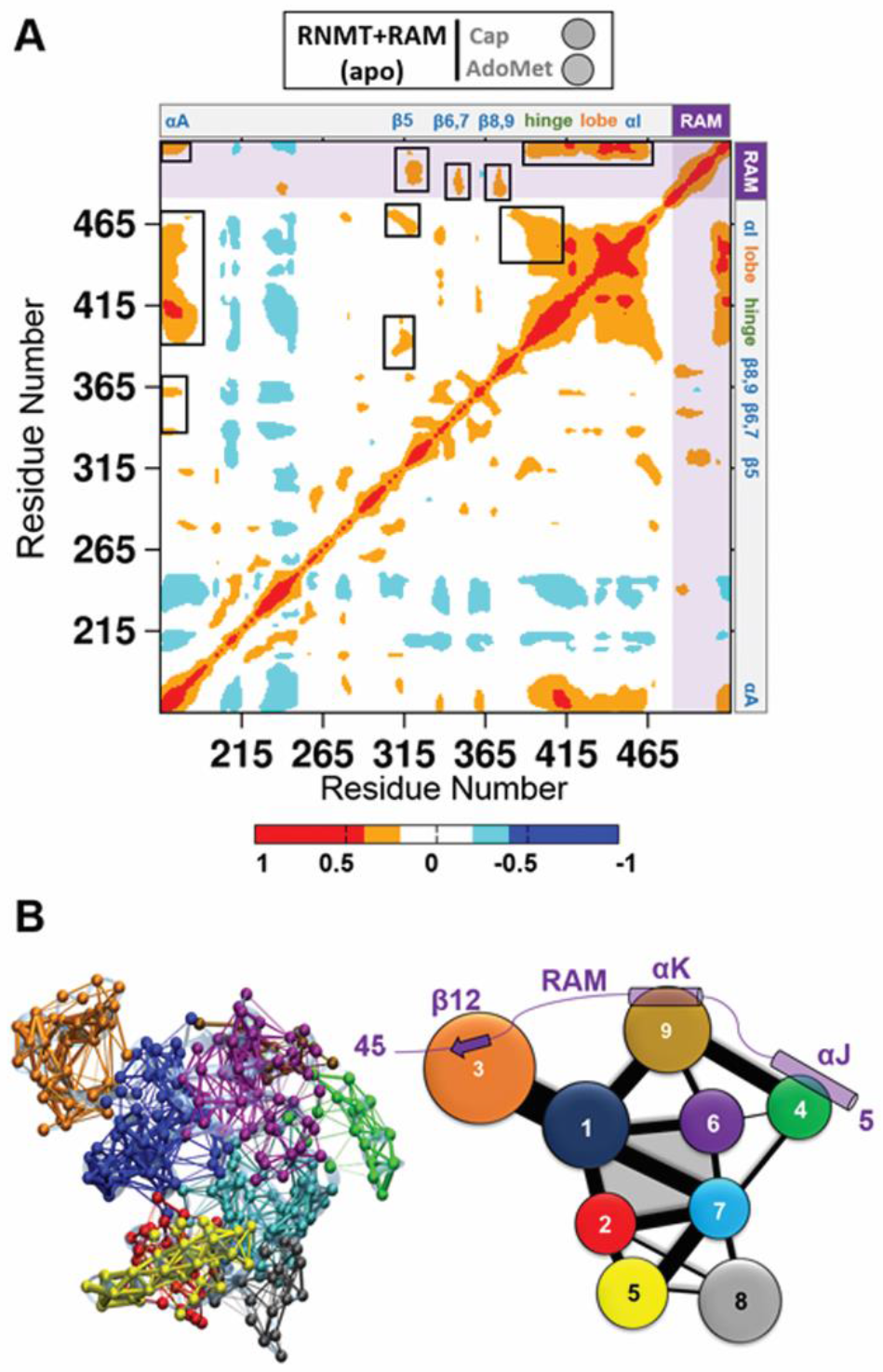
Allosteric networks in the RNMT-RAM complex. (A) Inter-residue cross correlation plot calculated for the *apo*-RNMT-RAM complex. Correlations range from −1 (anti-correlated, blue) to +1 (correlated, red). Positions of the key structural elements of RNMT are indicated at the top and right. RAM is included after the RNMT C-terminus and is highlighted by shading. Strong cross correlations that affect the RNMT active site are highlighted in black boxes in the upper triangle. (B) All-residue dynamic network and coarse-grained community analysis of the RNMT-RAM complex. *Left*: The all-residue dynamic network representation, with points (“nodes”) representing protein residues. The nodes are connected by the “edges” if they are found to be within 4.5 A of each other during the simulation. In addition, the nodes are coloured according to the communities they belong to. *Right*: Two-dimensional community diagram of RNMT-RAM. Each community is shown as a coloured circle, with a radius proportional to the size of the community (i.e. number of residues). The communities that were found to interact with each other are connected by edges, with the width of an edge proportional to intercommunity interaction strength. The active site is formed by the communities 1, 2, 6 and 7 and is shaded in grey.

Inter-residue correlation analysis of the *apo*-RNMT-RAM complex simulations revealed strong correlations between the key regions (Figure 7A). In particular, α-helix A was strongly correlated with the α-helix hinge and the lobe, and to a smaller extent with the β-strand lid. As expected, important correlated motions were also observed between the hinge and the lobe elements. Importantly, there is strong correlation between RAM and the structural elements forming the RNMT active site: in particular, the N-terminal region of RAM (residues 2-20) correlates with strands β5, β6-7 and β8-9, whereas the C-terminal region of RAM (residues 30-45) correlates with the hinge, the lobe, α-helix I and, most notably α-helix A. Hence, a vast matrix of correlated dynamic motions connecting the RNMT key regions with the N- and C-termini of RAM provided additional evidence of the allosteric modulation fulfilled by RAM.

The all-residue dynamic network and coarse-grained community network analyses (56, 60) were then performed to identify the allosteric paths for the RNMT regulation by RAM (Figure 7B). Both RNMT and RAM were included in the network analysis of the complex. The community assignment is in good agreement with the structure-based domain identification in RNMT. Four communities comprise the cap and AdoMet binding pockets: communities 1 and 2 include the upper and lower halves of helix αA, respectively; community 6 contains the β6-β9 lid region and helix αE, whereas community 7 includes the AdoMet binding Motifs I and II. These four communities displayed strong intercommunity connections, which is indicated by the width of the edges in the 2D community network diagram (Figure 7B). In addition, on the periphery, community 1 partially includes the lobe region and confirms the strong connections between helix αA and the lobe.

The network analysis also provided an evidence of the allosteric communication between the RNMT active site and RAM. First, community 1 (helix αA + lobe) showed strong connections with the C-terminal end of RAM (strand β12), which is included in community 3; second, the middle region of RAM (RAM helix αK, community 9) communicates with the active site communities 1 and 6; finally, helix J in the N-terminal region of RAM (RAM residues 2-15, community 4) showed strong connections with RAM helix K and also with communities 6 and 7 within the RNMT active site. The correlation and network analyses unveiled underlying long-range allosteric networks and paths that are crucial for allosteric regulation of RNMT by RAM.

### RAM residues 1-20 are critical for RNMT activation

The network analysis of the *apo*-RNMT-RAM complex revealed three key regions within RAM that communicate with the RNMT active site and are important for allosteric regulation (Figure 7B), namely a short strand β12 (in the RAM C-terminus; part of community 3) and helices αK (residues 24-30; part of community 9) and αJ (in the N-terminal end; part of community 4). The first two were known to provide a key stabilization effect on the lobe and the active site residues, as was shown in our recent computational and biochemical study (8), where we described in great detail the interactions of these regions with the RNMT lobe and α helix A, as well as explored an extensive set of *in silico* and *in vitro* mutants designed to test these interactions. On the other hand, the finding that a more remote N-terminal region of RAM (residues 1-20, containing helix αJ) also participates in the allosteric networks and contributes to regulating the RNMT activity was somewhat surprising and not immediately evident from the crystal structure. It was previously shown (61) that smaller fractions of the RNMT activation domain of RAM, e.g. RAM 1-35 and RAM 20-45, are sufficient to interact with and partially stabilize RNMT, however the entire domain (RAM 1-45) is required for maximal RNMT activation.

To provide further support to this finding, we designed an additional system in which the N-terminus of RAM was deleted, and simulated RNMT in complex with such a *truncated* form of RAM (residues 20-45). The conformational behaviour of RNMT in the cMD and aMD simulations was then analysed following the above described protocols. The simulations demonstrated that the truncated RAM maintained an overall stability, with the exception of the helix αK that exhibited much higher flexibility in the absence of the RAM N-terminus, which previously acted as an additional stabilizing anchor. In turn, RNMT largely demonstrated the same structure and dynamics as in the complex with a longer RAM (Figure 8A; compare with Figure 4B).

**Figure 8.**
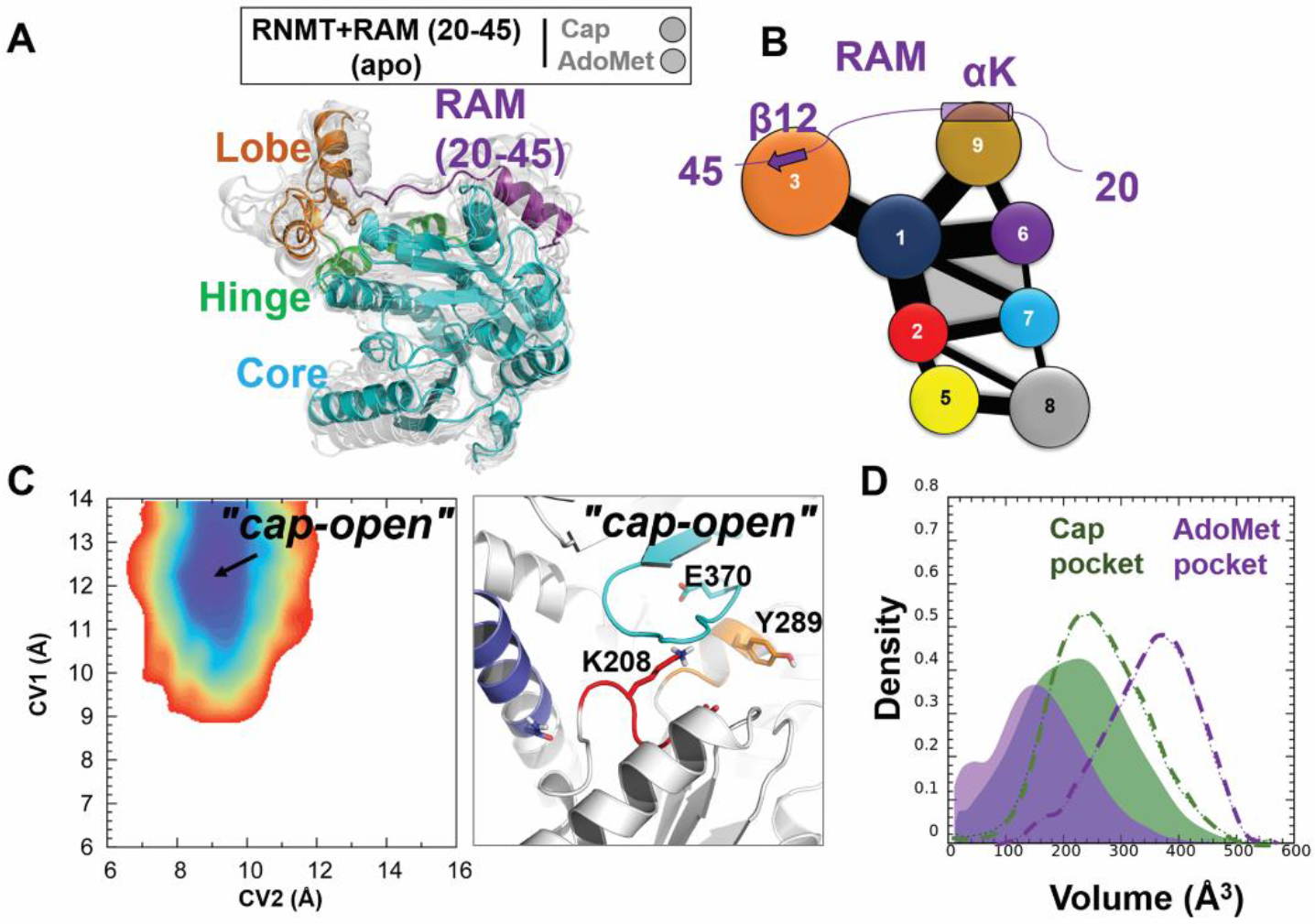
Modulation of the RNMT active site upon binding of a truncated form of RAM (20-45). (A) Conformational dynamics of RNMT in complex with the truncated RAM (20-45). The representative simulation snapshot is superimposed with multiple structures (light grey) that were sampled from the entire aMD trajectory and illustrate the range of conformational dynamics. The key structural elements are labelled and coloured as in Figure 4B. (B) The community network diagram of the RNMT-RAM (2-45) complex. The representation is the same as in Figure 7B. (C) *Left*: Free energy landscape (FEL) of *apo*-RNMT in complex with the truncated RAM. *Right*: The representative structure of the RNMT active site corresponding to the “*cap-open*” state. The representation is the same as in Figure 5. (D) Distribution of the volume of the cap (green) and AdoMet (magenta) binding pockets during the aMD simulation of *apo*-RNMT in complex with the truncated RAM. The representation is the same as in Figure 6.

As expected, the RNMT interactions with the C-terminal region of RAM were largely unaffected by the removal of the RAM N-terminus: the RNMT lobe was still well structured in the presence of the truncated RAM and contributed to maintaining the secondary structure elements of the hinge and α-helix A. However, a notable difference was observed for the RNMT β-lid: strands β8 and β9 were significantly destabilized, inducing a conformational change within the AdoMet binding pocket.

This dynamic behaviour was rationalized by performing the network analysis in the RNMT-RAM (20-45) system. The obtained community diagram (Figure 8B) is largely identical to the one with a full-length RAM (Figure 7B), with one important exception that the removal of the RAM helix J has led to the disappearance of community 4, which previously was interacting with communities 6 and 7 that comprise the AdoMet binding site. As a result, one component of the allosteric network between RAM and RNMT has been lost and the connectivity between RAM and the AdoMet binding site was significantly reduced.

This result is fully consistent with the free-energy landscape analysis performed for the simulation with the truncated RAM. The computed FEL of the active site of RNMT has yielded a qualitatively different picture (compare Figure 8C with Figure 5D): now there is only one free energy minimum, which corresponds to the “*cap-open*” state and has the same position and shape as in the case of the full-length RAM. This conformation of the active site (Figure 8C) favours binding of the cap; this is also confirmed by monitoring the volume of the cap pocket (Figure 8D), which remained largely the same as for RNMT with the full-length RAM (Figure 6C). In contrast, the previously observed second minimum, corresponding to the “*fully-open*” state in Figure 5D, has now completely disappeared, meaning that this conformation was not sampled at all when the N-terminal region of RAM was removed. The pocket volume analysis consistently showed that the AdoMet binding pocket has shrunk and cannot accommodate the substrate (Figure 8D). The representative snapshots provide a molecular explanation of such qualitatively different behaviour: the AdoMet binding pocket was occluded by the displacement of the strands β8-β9 and Motif I residues (Figure 8C); in particular, the sidechains of both K208 and Y289 have flipped to partially occupy the AdoMet binding site.

Altogether, our results provide an evidence of the importance of the RAM N-terminal region for the allosteric regulation of RNMT, and specifically its crucial role in controlling the AdoMet pocket through the allosteric network connecting the RAM N-terminus to the AdoMet binding site. The truncated RAM can still efficiently promote binding of the cap. However, in the absence of the N-terminal section of RAM, the AdoMet binding is precluded, which will affect the RNMT activity. This is consistent with the previous experimental observation (61) that a longer RAM 1-45 is required for maximal RNMT activation.

## DISCUSSION

In eukaryotes, nascent RNA polymerase II transcripts are cotranscriptionally modified at the 5’ end by the addition of the mRNA cap structure which is essential for an efficient transport, stability and translation of the mRNA (62-64). N7-methylation of cap guanosine is catalysed by the mammalian mRNA cap methyltransferase (RNMT), which is known to be activated by RAM (6, 62). Important efforts were carried out in an attempt to shed light into the molecular basis that could explain the allosteric modulation of RNMT activity induced by RAM including biophysical, crystallographic and modelling studies (8, 61, 65). The use of computer simulations and molecular modelling has shown to be of great value by providing molecular rationale for experimental research findings, offering a tool in situations when experimental methods could not succeed, and often guiding further experiments. In particular, molecular dynamics simulation method has grown into a robust tool for describing dynamics of biomolecular systems at atomic resolution. However, allosteric regulation involves conformational changes often taking place on the microsecond to millisecond timescales which are difficult to reach using standard molecular dynamics simulations (41, 58, 66, 67). In this work, we investigated RNMT activation by RAM by means of accelerated molecular dynamics (aMD) simulations, a popular enhanced sampling method. aMD simulations were able to sufficiently sample the free energy landscape of RNMT and to capture the conformational changes occurring within the RNMT active site that can account for the RAM-dependent activity of the enzyme.

To date, the lack of crystallographic data of mRNA methyltransferases with simultaneously bound substrates, AdoMet and cap, challenges the clarification of this methyl-transfer mechanism. Here, we have built a model of the human RNMT-RAM complex with the bound reaction substrates and validated it by using standard MD and aMD simulations. The cap fits into a pocket formed in-between α-helix A and strands β6-β9, which form the “lid” of the pocket, establishing polar and hydrophobic interactions with G_0_ of the cap. The polyphosphate chain is stabilized by a network of positively charged residues, namely R173, K180 and K208, elucidating the location of the nascent mRNA transcript bound to RNMT. The methyl donor AdoMet interacts with the conserved Motif I and Motif II loops; its adenine base is stacked between I228 and the cap-stabilizing residue Y289. The binding poses of the reaction substrates bound to RNMT-RAM were maintained throughout the unrestrained aMD trajectories, and the secondary structure conformations of key structural elements of RNMT (hinge and lobe) were essentially preserved. Interestingly, when AdoMet and the cap were simulated bound to RNMT in the absence of RAM, the original binding poses were equally retained despite the remarkable flexibility of the hinge, the lobe and α-helix A. Although we cannot reject the possibility that the substrates could eventually be destabilized on a longer time-scale, the results suggest that any stabilization effect of RAM on the RNMT active site mainly occurs prior to binding of the substrates.

We have explored possible mechanisms of such stabilization effect by building a comprehensive set of simulation systems (with/without substrates, with/without RAM) and performing standard and aMD simulations for them. The aMD simulations of *apo*-RNMT in the presence and absence of RAM have offered molecular and energetic insights into the RAM-induced conformational changes occurring in the RNMT active site. Free energy landscapes (FEL) representing the conformational space of the RNMT active site were computed for each simulated system. In the absence of RAM, the results suggest an equilibrium between a fully “closed” form of *apo*-RNMT (both substrate pockets are inaccessible) and a conformation in which binding of the cap is favoured. The “fully open” state, with optimal and accessible AdoMet and cap pockets, was not favoured in *apo*-RNMT. In contrast, the *apo*-RNMT-RAM complex has demonstrated an equilibrium between a “fully open” conformation of the enzyme’s active site and a conformation favouring binding of the cap, whereas the “closed” state was not sampled. This finding was further explained as a result of the stability observed in α-helix A and the β8-β9 lid that favours cap binding, along with the reorientation of Y289 and Motif I loop towards the cap binding pocket, which increases the accessibility to AdoMet. aMD simulations of RNMT-RAM with a single cap substrate showed a notably reduced barrier between the “fully-open” and the “cap-open” conformations, suggesting that the cap binding is likely to promote subsequent binding of AdoMet. This event was previously reported by our group (8) and can now be rationalized by a conformational shift of K208 towards the cap polyphosphate chain that favours the accessibility to AdoMet. It must be noted it remains possible that the induced fit mechanism might enable AdoMet or cap binding to RNMT in the absence of RAM, even when respective pockets deemed inaccessible. In summary, the studies showed that binding of RAM notably enhances selection of the RNMT active site conformations suitable for binding of substrates, providing a rationale to the increased RNMT catalytic activity measured in the presence of RAM.

Finally, the community network analysis performed on the aMD simulations of the RNMT-RAM complex has provided evidence for the long-distance allosteric communications between RAM and the active site residues. The study has confirmed strong correlations between RAM, lobe, hinge and α-helix A. The characterized allosteric networks have offered new evidences on putative interactions with helices αE, αD, strands β6-9 and Motif I, suggesting that the N-terminal region of RAM (residues 1-20) can directly affect the RNMT active site conformations. This hypothesis has been confirmed by FEL of the complex of RNMT with a truncated RAM (residues 20-40), which has failed to sample the conformations suitable for AdoMet binding or the “fully open” state. This result is in agreement with previous studies that showed that a truncated form of RAM (20-45) was able to bind to RNMT but failed in the activation process of the enzyme (6).

In summary, aMD simulations have provided new insights into the molecular mechanisms of the RNMT allosteric regulation and have shed light on the conformational changes induced by RAM on the RNMT active site favouring binding of the reaction substrates. We have identified a number of residues critical for binding of both substrates and RAM; some of these have not been previously tested by site-directed mutagenesis experiments. Such information is critical for drug discovery efforts aiming to reduce the cap methyltransferase activity often overexpressed in tumour cells, and that will benefit of a clearer insight into the mechanism of action of RNMT-RAM complex. Moreover, the recent crystallographic structures of methyltransferases from different flavivirus such as Zika (69) constitute a very promising starting point for understanding the molecular details of the methyltransferase machinery in viruses and will offer opportunities for structure-based rational design of antiviral inhibitors.

## Supporting information

AdoMet_cap_binding_mode

## SUPPLEMENTARY FIGURES AND TABLES

**Supplementary Figure S1.**
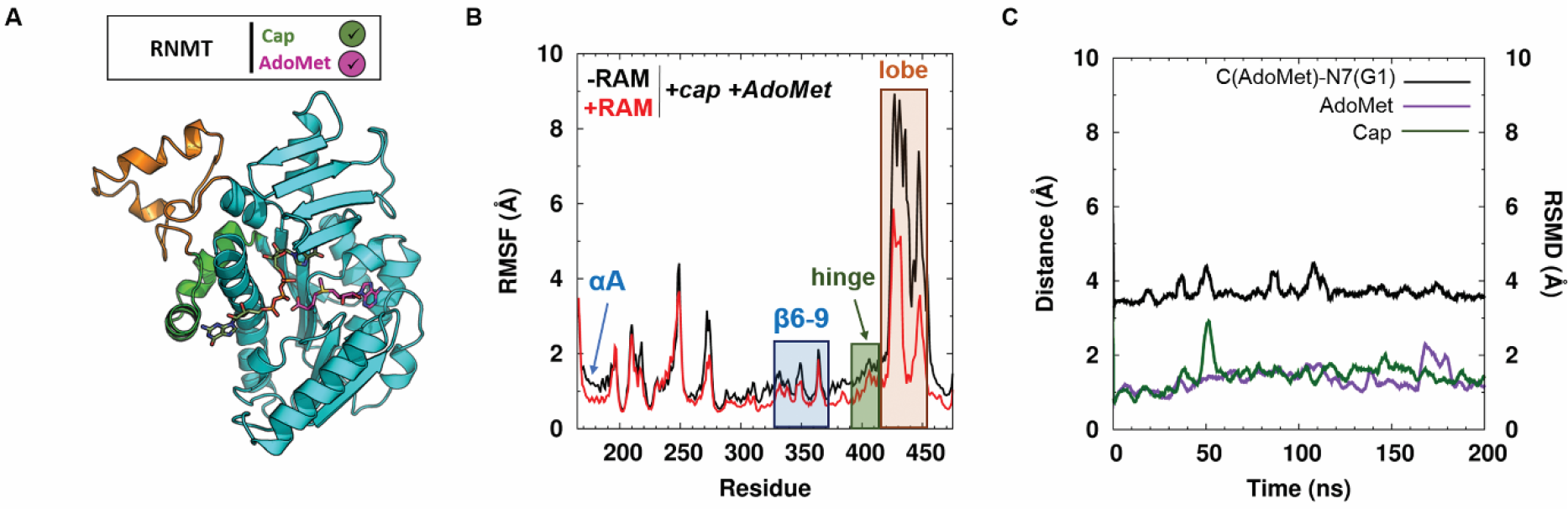
RNMT modelled with the bound cap and AdoMet in the absence of RAM. (A) Representative snapshot of RNMT in complex with the cap (green) and AdoMet (magenta) in a methyl-transfer reactive conformation. (B) RMSF values (in Å) of the RNMT obtained from the aMD simulation. (C) Time-evolution of the distance between AdoMet (CH3) and the cap G_0_ (N7 atom) (black line). The RMSD of AdoMet (magenta) and cap (green) are plotted against the secondary (right) vertical axis.

**Supplementary Table S1.**
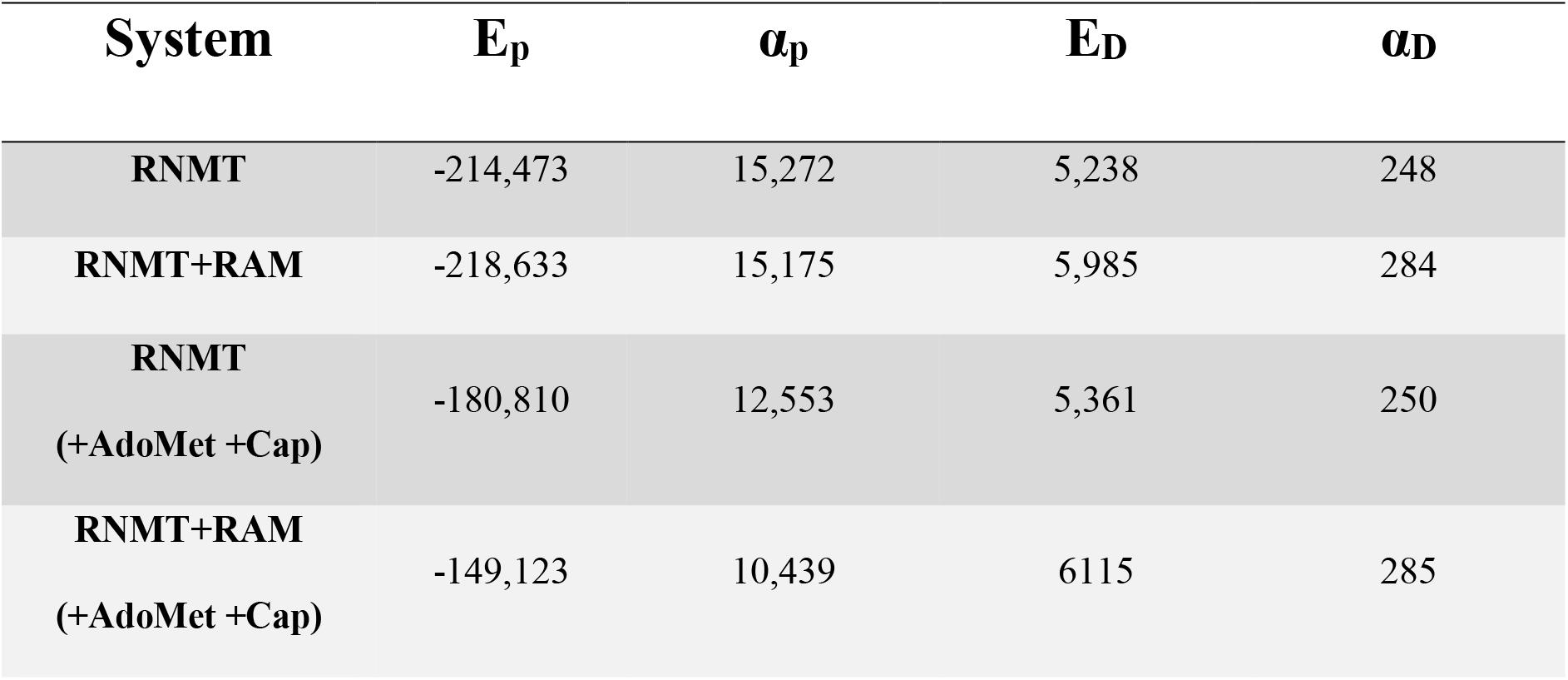
aMD parameters used in the present simulations. The choice of the parameters was initially based on the guidelines from previous works (43-45). Further adjustments were done using the preliminary aMD simulations. Values of all the parameters are given in kcal mol^−1^.

| **System** | **E<sub>p</sub>** | **α<sub>p</sub>** | **E<sub>D</sub>** | **α<sub>D</sub>** |
| --- | --- | --- | --- | --- |
| **RNMT** | -214,473 | 15,272 | 5,238 | 248 |
| **RNMT+RAM** | -218,633 | 15,175 | 5,985 | 284 |
| **RNMT**<br>**(+AdoMet +Cap)** | -180,810 | 12,553 | 5,361 | 250 |
| **RNMT+RAM**<br>**(+AdoMet +Cap)** | -149,123 | 10,439 | 6115 | 285 |

**Supplementary Table S2.** Key RNMT-cap and RNMT-AdoMet hydrogen-bonding and salt-bridge interactions observed in the simulations. The average distances for the key interactions computed over the cMD and aMD trajectories are given in Å, and the standard errors are reported in parentheses. Atoms between which the distance is measured are listed in the left two columns.

| Interaction |  | Distance (Å) |  |
| --- | --- | --- | --- |
| RNMT | Cap | cMD | aMD |
| N176 (ND2) | (O3') | 4.0 (0.5) | 4.3 (1.0) |
| H288(ND2) | (O6) | 3.9 (0.7) | 4.1 (1.2) |
| Y289 (OH) | (O6) | 2.9 (0.4) | 3.7 (1.5) |
| E370 (OE2) | (N1) | 2.8 (0.1) | 2.9 (0.4) |
| E370 (OE1) | (N2) | 3.0 (0.2) | 3.0 (0.4) |
| Y467 (OH) | (N3) | 3.1 (0.4) | 4.2 (1.4) |
| Y467 (OH) | O2') | 3.3 (0.5) | 3.6 (0.9) |
| RNMT | AdoMet | cMD | aMD |
| K180 (NZ) | (COO<sup>-</sup>) | 2.8 (0.1) | 2.9 (0.3) |
| G205 (CO) | (NH<sub>3</sub><sup>+</sup>) | 2.8 (0.1) | 2.8 (0.2) |
| D227 (OD1) | (O2/O3) | 2.5 (0.1) | 2.6 (0.1) |
| D261 (OD1) | (NH2) | 2.8 (0.1) | 2.8 (0.1) |
| S262 (NH) | N1 | 3.2 (0.2) | 3.2 (0.2) |
| Q284 (CO) | (NH<sub>3</sub><sup>+</sup>) | 2.8 (0.1) | 2.9 (0.3) |

